# Mono-homologous linear DNA recombination by the non-homologous end-joining pathway as a novel and simple gene inactivation method: a proof of concept study in *Dietzia* sp. DQ12-45-1b

**DOI:** 10.1101/300723

**Authors:** Shelian Lu, Yong Nie, Meng Wang, Hong-Xiu Xu, Dong-Ling Ma, Jie-Liang Liang, Xiao-Lei Wu

## Abstract

Non-homologous end-joining (NHEJ) is critical for genome stability because of its roles in double-strand break repair. Ku and ligase D (LigD) are the crucial proteins in this process, and strains expressing Ku and LigD can cyclize linear DNA *in vivo.* Herein, we established a proof-of-concept mono-homologous linear DNA recombination for gene inactivation or genome editing by which cyclization of linear DNA *in vivo* by NHEJ could be used to generate non-replicable circular DNA and could allow allelic exchanges between the circular DNA and the chromosome. We achieved this approach in *Dietzia* sp. DQ12-45-1b, which expresses Ku and LigD homologs and presents NHEJ activity. By transforming the strain with a linear DNA mono homolog to the sequence in chromosome, we mutated the genome. This method did not require the screening of suitable plasmids and was easy and time-effective. Bioinformatic analysis showed that more than 20% prokaryotic organisms contain Ku and LigD, suggesting the wide distribution of NHEJ activities. Moreover, the *Escherichia coli* strain also showed NHEJ activity when the Ku and LigD of *Dietzia* sp. DQ12-45-1b were introduced and expressed in it. Therefore, this method may be a widely applicable genome editing tool for diverse prokaryotic organisms, especially for non-model microorganisms.

**IMPORTANCE:** The non-model gram-positive bacteria lack efficient genetic manipulation systems, but they express genes encoding Ku and LigD. The NHEJ pathway in *Dietzia* sp. DQ12-45-1b was evaluated and was used to successfully knockout eleven genes in the genome. Since bioinformatic studies revealed that the putative genes encoding Ku and LigD ubiquitously exist in phylogenetically diverse bacteria and archaea, the mono-homologous linear DNA recombination by the NHEJ pathway could be a potentially applicable genetic manipulation method for diverse non-model prokaryotic organisms.

## INTRODUCTION

Bacterial genome editing such as deletion, mutation, and insertion of genes is an efficient method to understand gene functions and modify metabolic activities. The most often used gene recombination methods in bacteria include the bacteriophage recombination system using the linear double-strand DNA (dsDNA) as recombination substrate (1–3), the single or double crossover homologous recombination (HR) using the circular plasmid DNA as substrate (4, 5), and the transposase-mediated transposition recombination (6, 7). Recently, several genome editing technologies have emerged, which are mediated by the targeted nucleases, including zinc finger nucleases (ZFNs) (8), transcription activator–like effector nucleases (TALENs) (9), and short palindromic repeats (CRISPR)-associated Cas9 endonuclease (10). All these methods generally require suitable plasmids and a laborious process such as cloning of genes into plasmids. Moreover, it is nearly impossible to find compatible plasmids for diverse prokaryotic organisms in nature, especially for the non-model microorganisms. Therefore, developing efficient and suitable genetic manipulation methods is still of great necessity.

Generally, a genetic manipulation method is developed according to a natural biological process. For example, the HR method was developed following a natural HR process to repair DNA double-strand break (DSB) (11), which evolves in all cellular organisms (12). Another DSB repairing process is the non-homologous end joining (NHEJ) process, which was first discovered in mammalian cells (13). During the eukaryotic NHEJ process, the ends of broken DNA are approximated by the DNA-end-binding protein Ku (Yku70/Yku80 in yeast) and then joined by an ATP-dependent DNA ligase IV/XRCC4/XLF (LXX) complex (LigD, Dnl4/Lif1/Nej1 in yeast) (13).

Consequently, a homologous DNA template is not needed (13, 14). Recently, NHEJ was also discovered in prokaryotes with the evidence of the circulation of a linear plasmid DNA in Ku and LigD containing mycobacteria (15, 16) and the circulation of a linear DNA in *Escherichia coli* expressing mycobacterial Ku and LigD (17). In addition, the Ku homodimer and LigD are crucial proteins in the bacterial NHEJ process (18–20), whose presence could indicate NHEJ activities in bacteria (21–23). NHEJ also offer protection to bacteria when only a single copy of the genome is available such as after sporulation or during stationary phase under environmental stresses (24, 25).

Similar to the HR DSB repairing process, the NHEJ process may also be used for genetic manipulation. In this case, a target homologous sequence can be amplified, PCR fused with a selective marker to form a linear DNA fragment and then transformed into the recipient bacterium, which expresses Ku and LigD. If the linear DNA can be cyclized by the NHEJ pathway *in vivo*, the resultant circular DNA may act as a circular incompatible plasmid in bacterial genetic manipulation (26). Because the linear/circular DNA without replication region cannot be replicated, the transformed cells surviving against selective stresses such as antibiotic should be mutants with insertion in the target sequences. Consequently, the target gene can be knocked out by using the NHEJ pathway.

Herein, we proposed a novel and much simpler gene knockout method by using the NHEJ pathway in *Dietzia* sp. DQ12-45-1b. *Dietzia* strains found in diverse environments are powerful *n*-alkane degraders and potential pathogens. They can utilize a wide range of compounds as the sole carbon source such as hydrocarbons and aniline (27–29). Moreover, a number of *Dietzia* strains can be used as potential probiotics to inhibit fecal *Mycobacterium avium* subspecies *paratuberculosis in vitro* (30) and might be useful for the treatment of patients with Crohn’s disease (31, 32). They are non-model gram-positive bacteria and lack efficient genetic manipulation systems, but they express genes encoding Ku and LigD. The NHEJ pathway in *Dietzia* sp. DQ12-45-1b was evaluated and was used to successfully knockout eleven genes in the genome. Since bioinformatic studies revealed that the putative genes encoding Ku and LigD ubiquitously exist in phylogenetically diverse bacteria and archaea, the mono-homologous linear DNA recombination by the NHEJ pathway could be a potentially applicable genetic manipulation method for diverse non-model prokaryotic organisms.

## MATERIALS AND METHODS

### Bacterial strains, plasmids, media, and primers

The bacterial strains and plasmids used in this study are listed in Table 1. All primers are listed in Table S1. *Dietzia* sp. DQ12-45-1b, isolated from oil-production water, has been intensively studied in our laboratory (28, 33-38). *Dietzia* sp. DQ12-45-1b was cultured in GPY medium (1% [wt/vol] glucose, 1% [wt/vol] tryptone, 0.5% [wt/vol] yeast extract) at 30°C on a rotary shaker at 150 rpm. The alkane hydroxylase-rubredoxin fusion gene (*alkW1*) (34) mutant of *Dietzia* sp., DQ12-45-1b (*alkW1*^-^), was grown in GPY medium with 40 mg/liter kanamycin. To determine their functions in alkane degradation, strains *alkW1*^-^ and *Dietzia* sp. DQ12-45-1b were grown in minimal medium (4) supplemented with 0.1% (vol/vol) hexadecane as the sole carbon source at 30°C, as described previously (34). *E. coli* DH5α was grown in lysogeny broth (LB) medium at 37°C. *E. coli* strains and *Dietzia* strains harboring plasmids were grown with appropriate antibiotics (ampicillin, 100 μg/ml; kanamycin, 30 μg/ml; streptomycin, 30 μg/ml).

**TABLE 1.**
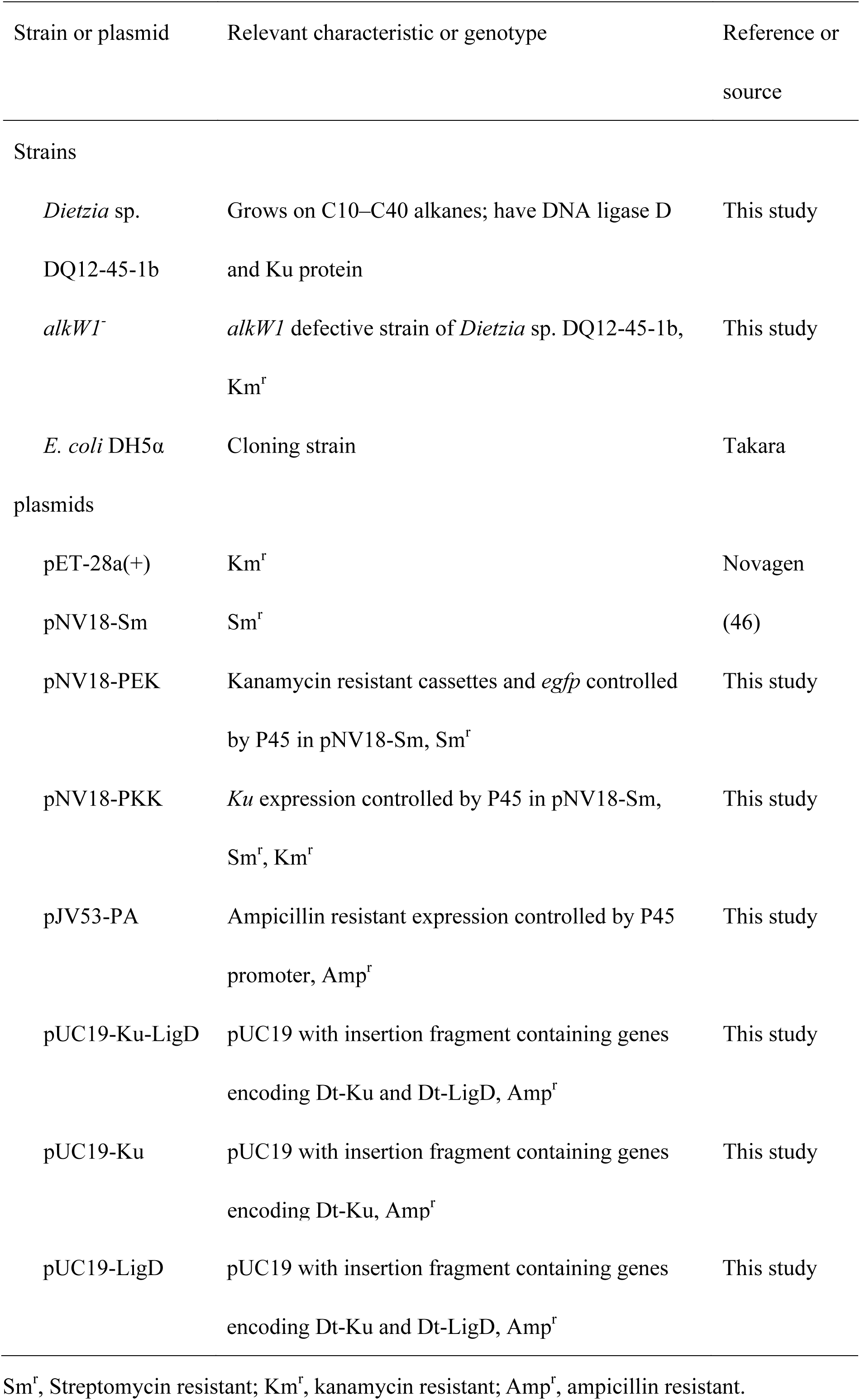
List of strains and plasmids used in the study

The vector pNV18-PEK (Table 1) was constructed as follows. Firstly, the promoter p45 DNA fragment was amplified from plasmid pNV18-Dsred (44) with primer p45-EF and p45-BR, *egfp* gene from pK18-egfp (unpublished result) with primer egfp-BF and egfp-HR, and kanamycin-resistant gene (Km) from plasmid pK18 (39) with primer Km-HF and Km-PR. Then the promoter p45 was clone into pXL1801 (40) with EcoRI and BamHI restriction sites, *egfp* with BamHI and HindIII restriction sites and Km with HindIII and PstI restriction sites sequentially, obtaining the plasmid pNV18-PEK. The plasmid pJV53-PA was also constructed. Briefly, a large fragment of vector pJV53 was first amplified with primer 53-HF and 53-ER (Table S1). The resulting 3,013 bp linear fragment lacking the acetamidase promoter and Chec9c 60-61 was digested with EcoRI and HindIII. The promoter p45 was amplified from pNV18-PEK with primer p45-EF and p45-(amp)-R (Table S1), and the ampicillin resistant gene was amplified from pUC19 with primers amp-(p45)-F and amp-HR (Table S1). The p45 and ampicillin resistant gene was fused by PCR with primer p45-EF and amp-HR, resulting in PA DNA fragment. Then PA was ligated into linear pJV53 by the EcoRI and HindIII restriction sites to obtain the plasmid pJV53-PA.

### Sequence analysis

Genes encoding the Ku and LigD homologs in the genome of *Dietzia* sp. DQ12-45-1b, designated as Dt-Ku and Dt-LigD, were identified by comparing the proteome of strain DQ12-45-1b (unpublished result) against the non-redundant (NR) database of protein sequences at the National Center for Biotechnology Information (NCBI) using BLASTP (41). The conserved domains of proteins were determined by comparing the protein sequences against the conserved domain database (CDD) at NCBI (42). To identify the Ku and LigD homologous genes in prokaryotic organisms, we searched the KEGG Orthology (KO) (43) for ID K10979 and K01971, which indicated Ku and LigD, respectively, in the Integrated Microbial Genomes (IMG) system (44) against a total of 25,270 bacterial genomes and 528 archaeal genomes (until April 2015). Sequence alignment and phylogenetic analysis were performed in MEGA6 (45).

### NHEJ assay in *Dietzia* sp. DQ12-45-1b

To identify the circular efficiency of different linear DNA ends, three types linear DNA was generated. The linear DNA fragment with 5′-“A” tails (LA18) was amplified from 4 ng plasmid pNV18-Sm (46) using LA Taq (Takara, Tokyo, Japan) under the conditions suggested by the manufacturer with the primers pNV18-F and pNV18-R, followed by treatment with DpnI digestion (Takara, Tokyo, Japan) for 4 h at 37°C to eliminate the circular plasmid pNV18-Sm template. Secondly, the plasmid pNV18-Sm was digested by HindIII to produce 5′-overhang DSB fragments HD18, and by HindIII and BamHI simultaneously to produce 5′-overhang DSB fragments HB18 overnight, respectively (Fig. 1). All three linear DNA fragments contained intact streptomycin-resistant gene (Fig. S1). Gel purification was performed to remove the undigested plasmids and collect the linear plasmid only with DNA purification kit (Tiangen biotech, Beijing, China). Five microliter purified linear DNA fragments, LA18 (108 ng), HD18 (437 ng), and HB18 (188 ng), were then transformed into 100 μL competent cells of strain DQ12-45-1b (~10^10^ CFU/ml and the transformation efficiency was 10^6^ CFU/μg DNA), respectively. After transformation, cells were recovered at 30°C for 4 h, followed by plating on LB agar containing streptomycin. Positive clones were numbered and picked after 3 days incubation. The method of competent cell preparation and electro-transformation was as previously described (33).

**FIG 1.**
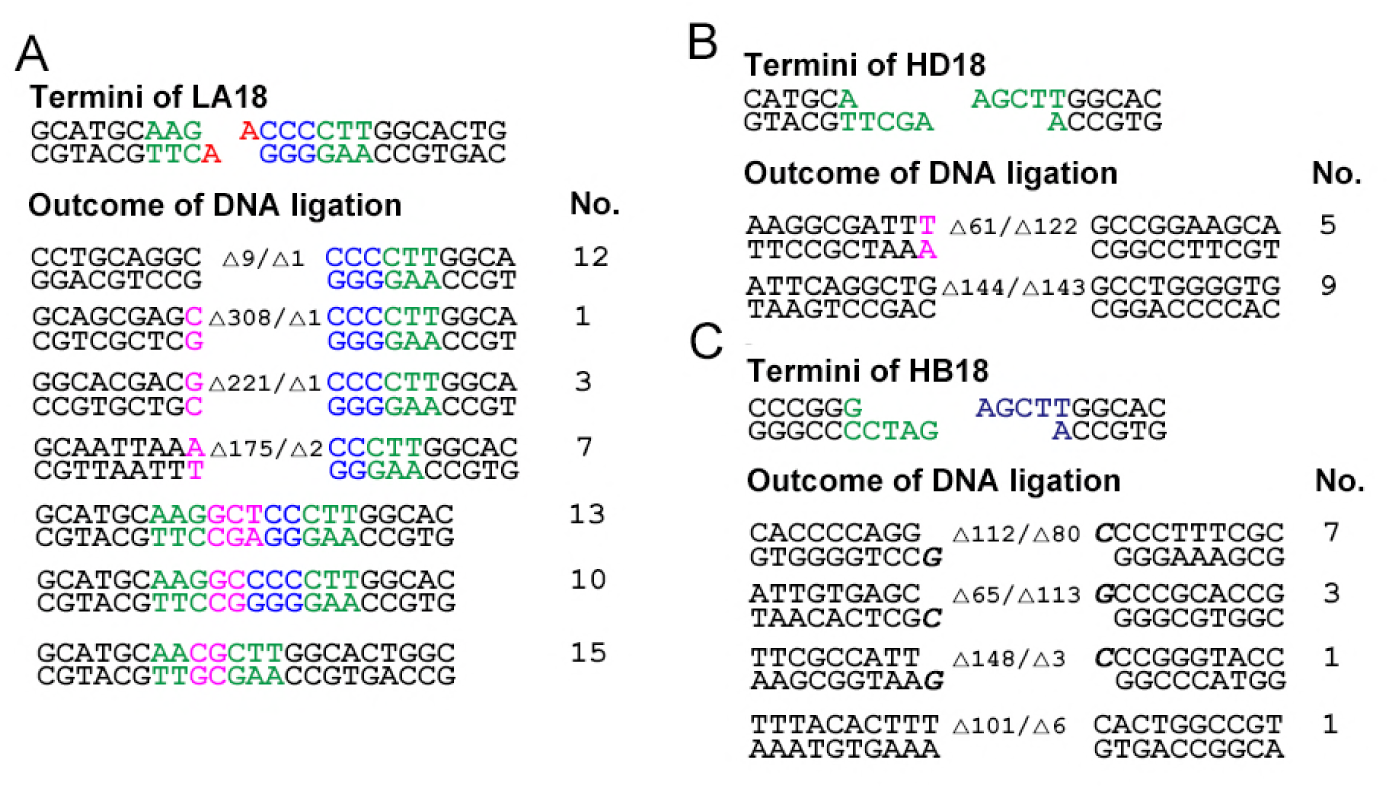
Sequences of DSB junction in LA18 (A), HD18 (B), and HB18 (C). The 5′-“A” tail is red. Hindlll and BamHI digestion sites are shown in green and purple, respectively. “CCC” label in LA18 added by PCR is in blue. The non-templated inserted nucleotides are in pink and the number of deleted nucleotides is shown at the junction site.

The resultant streptomycin-resistant colonies on LB agar were then suspended in 10 μL double distilled water and incubated at 98°C for 30 min to release DNA from the cells, followed by centrifugation at 20,000 ×*g* for 5 min. The DNA-containing supernatant was then subjected to PCR using r*Taq* (Takara, Tokyo, Japan) for 26 cycles with the primers IdenF and IdenR to amplify the joint region of the circular DNA. The PCR product was purified, cloned into pGEM-T (Promega, Madison, WI, USA). Recombinant plasmids were transformed into *E. coli* DH5α for white-blue screening, and the white colonies were sequenced to assess NHEJ activities.

To determine the role of *ku* gene in *Dietzia* sp. DQ12-45-1b, we constructed a *ku* inserted mutant with streptomycin resistance gene using a double-homologous-recombination method as described previously (Fig. S2) (47–49) and the primers used were listed in Table S1. The Ku expression vector for complementation studies were constructed as follows: the *ku* gene was amplified from *Dietzia* sp. DQ12-45-1b chromosomal DNA with the primers Ku-BF and Ku-HR (Table S1), and then the 1046 bp DNA fragment was ligated into pNV18-PEK using BamHI and HindIII restriction sites to construct a new plasmid, pNV18-PKK (Table 1). Vector pNV18-PKK was introduced into *ku* mutant cells via electroporation as described previously to obtain the complementary mutant cell (33). The effect of *ku* gene on the circular function of linear DNA in *Dietzia* sp. DQ12-45-1b were identified on the *ku*-complementary mutant cell using the linear plasmid DNA pJV53-PA (L53) with a p45 promoter and an ampicillin resistance gene (Fig. S3), which was amplified from plasmid pJV53-PA with the primers p45-F and 53-ER (Table S1) followed by being processed with DpnI digestion (Takara, Tokyo, Japan) for 4 h at 37°C to eliminate the circular plasmid pNV18-Sm template before being transformed into cells as described above, by taking the treatments operated on the WT cells and *ku* mutant cells transformed with pNV18-PEK as the negative controls (Table 1). After the 3-day incubation, ampicillin resistance colonies were counted as described above.

### Expression of Dt-Ku and Dt-LigD and NHEJ assay in *E. coli*

The genes coding for Dt-Ku protein and Dt-LigD were amplified from the genome of strain DQ12-45-1b using PrimeSTAR HS DNA Polymerase (Takara, Tokyo, Japan) according to the manufacturer’s instructions with the primers, KuF and KuR for Dt-Ku and LigDF and LigDR for Dt-LigD. The PCR products were purified and digested with the KpnI and EcoRI for Dt-Ku and with HindIII and KpnI for Dt-LigD. The plasmid pUC19-Ku and pUC19-LigD were then constructed by cloning *ku* and *ligD* DNA sequences into the corresponding sites of the plasmid pUC19. Similarly, the plasmid pUC19-Ku-LigD was constructed by cloning these two fragments into the HindIII and EcoRI sites of the plasmid pUC19, using standard methods (50), and then transformed into *E. coli* DH5α to construct the strain expressing single Dt-Ku or Dt-LigD and both of them, respectively.

To assess Dt-LigD and Dt-Ku activities, a heterogeneous expression test was performed. A linear DNA fragment P28 (3,818 bp, containing the pBR322 origin and kanamycin-resistant gene) was amplified from pET-28a(+) (Qiagen, Hilden, Germany) with the primers pET-28F and pET-28R. Specifically, 4 ng pET-28a(+) was used as template for PCR with PrimeSTAR HS DNA Polymerase (Takara, Tokyo, Japan) according to the manufacturer’s instructions. Then the PCR product was treated with DpnI digestion (Takara, Tokyo, Japan) for 4 h at 37°C to degrade the circular plasmid DNA and only the 3.8 kb band was collected for further purification with DNA purification kit (Tiangen biotech, Beijing, China). The pUC19, pUC19-Ku, PUC19-LigD and pUC19-Ku-LigD (100 ng) were then transformed by 42 °C heat shock for 90 s into 100 μL *E.coli* DH5α. After transformation, 900 μL of room temperature SOC medium was added to the tubes, and the cells were allowed to recover at 37°C with shaking at 300 rpm for one hour. 50 μL cells were cultured on LB agar containing 100 μg/ml ampicillin. The bacteria expressing no target protein or expressing Ku, LigD and Ku-LigD were prepared competent cells and then electro-transformed fragment P28 (500 ng) according to the standard instructions (50). All the cells were then collected by centrifugation (1,500 ×*g*, 5 min) and cultured on LB agar containing 100 μg/ml ampicillin and 40 μg/ml kanamycin. The transformation of pUC19 and P28 into *E. coli* DH5α was used as a negative control. The kanamycin resistant colonies indicated that the P28 was cyclized, allowing the expression of the kanamycin-resistant gene. Plasmids were then extracted from the positive colonies and digested with NdeI, followed by electrophoresis to determine the lengths of digested fragments.

### Gene disruption in *Dietzia* sp. DQ12-45-1b using mono-homologous arm linear DNA fragment

We used the *alkW1* gene as an example to determine whether NHEJ could be used for target gene manipulation. First, the homologous DNA fragment (from 143^rd^ to 552^nd^*nt* of *alkW1* gene), was amplified from the genome of strain DQ12-45-1b using primers 143-F and 552-R. For easy identification, a triplet cytidine (CCC) tag was additionally designed at the 5′ end of primer 552-R. Secondly, the kanamycin resistance cassette P45-EGFP-Km with a P45 promoter and an *egfp* gene as the reporter was amplified from the plasmid pNV18-PEK using the primer P45-F and Km-. Finally, the homologous DNA and P45-EGFP-Km cassette were fused and amplified by fusion PCR (51) using the primer P45-F and 552-R. The generated fragment (alk-Km) with homologous DNA of *alkW1* and kanamycin resistance cassette was purified and 5 μl of the purified product (500 ng) transformed into *Dietzia* sp. DQ12-45-1b as previously described (33). Cells were then grown on LB plate with kanamycin for 4 days. The kanamycin-resistant colonies were subjected to PCR amplification of gene *alkW1* with the primers alkW1-F and alkW1-R, which was further verified by sequencing. The recombinant efficiency was expressed as the number recombinants per microgram DNA divided by the cell competency which was indicated by the transformation efficiency with plasmid pNV18-Sm. The expression of AlkW1 in the cells cultured in liquid GPY medium or minimal medium with hexadecane as the sole carbon source was detected by western blot analysis as described previously (40).

To verify whether this method could be used for other genes, we selected seven genes encoding histidine kinase of two-component system and two other genes (Table 2) for disruption. The homologous linear DNA fragments were constructed and transformed as described above. The kanamycin-resistant colonies were selected and verified by PCR, fluorescence microscopic analysis and sequencing.

**TABLE 2.**
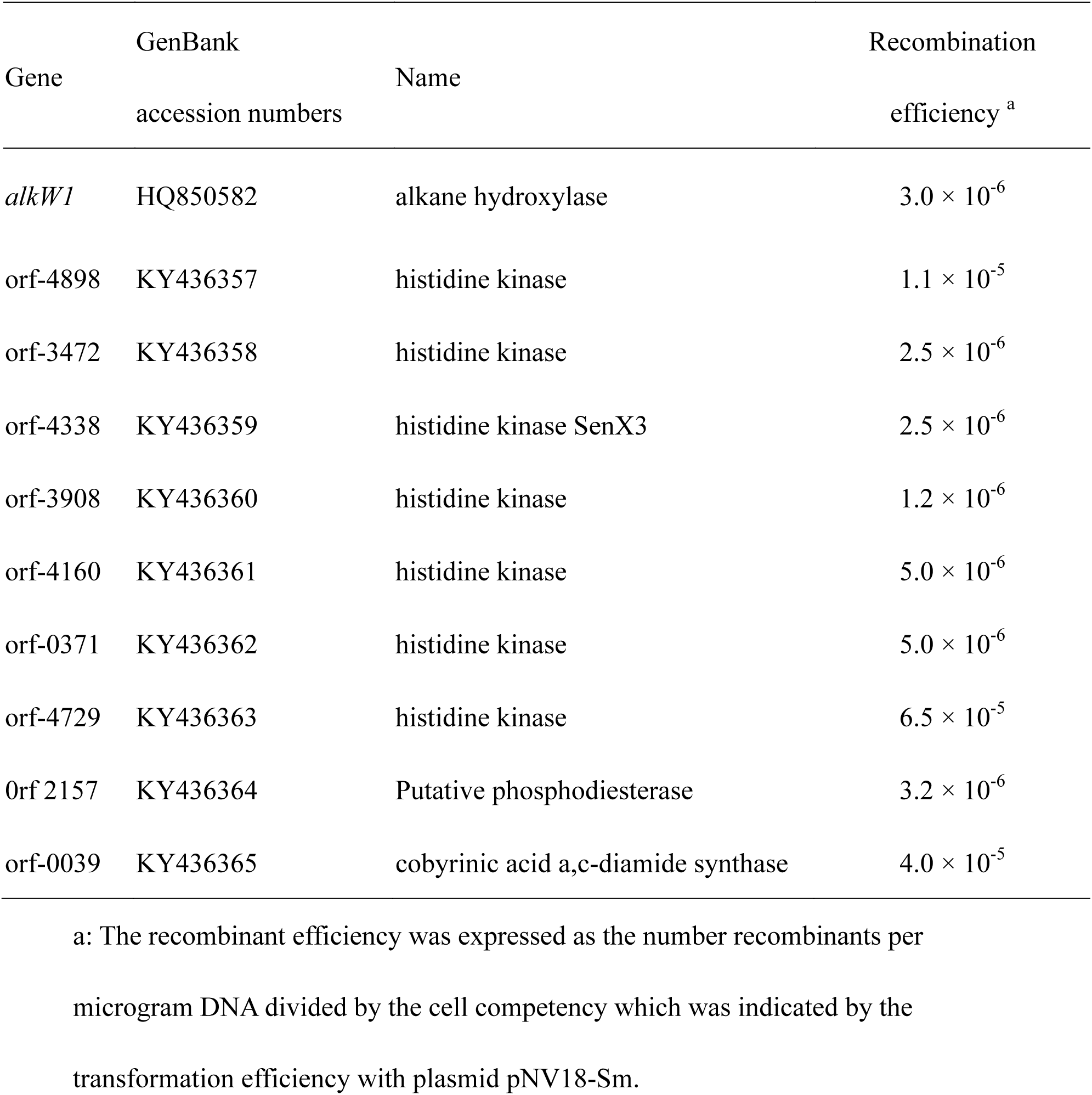
List of genes mutant in *Dietzia* sp. DQ12-45-1b

### Nucleotide sequence accession number

The GenBank accession numbers of Dt-Ku and Dt-LigD in *Dietzia* sp. DQ12-45-1b were KP074897 and KP074898, respectively. The GenBank accession numbers of the seven histidine kinases, putative phosphodiesterase, and cobyrinic acid a,c-diamide synthase were listed in Table 2.

## RESULTS

### NHEJ activity in *Dietzia* sp. DQ12-45-1b and *E. coli* expressing Ku and LigD

Genes encoding Ku and LigD homologs, Dt-Ku and Dt-LigD, were identified by screening the genome of *Dietzia* sp. DQ12-45-1b. These homologs showed 57% and 44% amino acid identities with Mt-Ku and Mt-Lig in *Mycobacterium* (18–20), respectively. To evaluate their NHEJ activities in strain DQ12-45-1b *in vivo*, the linear DNA fragments, LA18 with short overhang ends, HD18 with long overhang complementary ends, and HB18 with long overhang non-complementary ends (Fig. 1), were transformed into the strain DQ12-45-1b. In addition, negative controls were used in which the LA18, HD18, and HB18 fragments were transformed into *E. coli* DH5α that did not express components of the NHEJ pathway. No streptomycin-resistant colonies were detected in the negative controls, while transformation of LA18, HD18, and HB18 fragments into strain DQ12-45-1b generated 2130, 359, and 229 streptomycin-resistant colonies per microgram of linear DNA, respectively, indicating that the streptomycin-resistant gene originally embedded in the linear DNA fragments was functional in strain DQ12-45-1b (Fig. S4). Since the streptomycin gene containing DNA could be replicated only if it was in circular form, these results suggested that the linear DNA fragments were successfully cyclized in the strain, indicating that the NHEJ pathway was active *in vivo* in strain DQ12-45-1b. Among the three linear DNA fragments, it was more difficult for linear DNA with longer 5′-overhang ends to cyclize.

In addition, the three linear DNA fragments were cyclized differently by NHEJ activity. In case of LA18 with short overhang ends, 61 positive colonies were sequenced and 7 different NHEJ joints were detected (Fig. 1A). The overhung “A” at the ends was deleted before ligation in all colonies, suggesting the presence of a 5′-3′ exonuclease activity in strain DQ12-45-1b. Non-templated insertions and sequence deletions were detected in 38 and 23 out of the 61 colonies, respectively. Among the 38 colonies (62% of the 61 colonies) with insertion, three types of non-templated insertion sequences were identified. Two of them were detected in 25 colonies composed of G+C nucleotides (Fig. 1A). Among the 23 colonies with sequence deletions, 12 colonies contained short-scale (9 bp) deletions and the remaining 11 colonies contained 175–308 bp long deletions. It is notable that the deletions, being short or long, happened at the ends of the side without “CCC” tag, suggesting that the repeated G+C sequence might protect the linear DNA from exonuclease activity. In contrast, for HD18 and HB18 with long overhang ends, long unidirectional deletions up to 148 bp, were detected at both ends (Fig. 1B and 1C). The results further confirmed the DNA polymerase and exonuclease activities in the NHEJ process in strain DQ12-45-1b (15).

To evaluate the NHEJ activities of Ku and LigD in strain DQ12-45-1b, *ku* mutant strain was constructed. It was notable that *ligD* mutant was failed to be obtained even though two different methods had been tried to disrupt ligD gene from the strain (data not shown), The result suggested that *ligD* might be an essential gene for the strain DQ12-45-1b. The linear DNA L53 with blunt ends was transformed into wild type, *ku* gene mutant, and the mutant with *ku* complementation to observe the NHEJ activities. No ampicillin-resistant colonies were obtained from the *ku* mutant cells, while there were 463 and 351 ampicillin-resistant colonies per microgram of linear DNA in the wild type cells and the mutant with *ku* complementation (Fig. S5), indicating that the linear DNA L53 were cyclized in the cells. In addition, the L53 fragment was also transformed into *E. coli* DH5α, and no ampicillin-resistant colonies were detected, suggesting that there was no circular plasmid contaminant.

To further confirm the cyclization capability of Dt-Ku and Dt-LigD, we transformed pUC19, pUC19-Ku, pUC19-LigD and pUC19-Ku-LigD firstly into *E. coli* DH5α following electro-transformation P28 and successfully recovered colonies in presence of kanamycin, while no kanamycin-resistant colony was detected in the bacteria including pUC19, pUC19-Ku and pUC19-LigD. From the kanamycin-resistant colonies, plasmids were extracted and digested by NdeI to further verify the circulation results. Three bands with the expected sizes of about 1.2 kb, 5.2 kb, and 3.8 kb were observed, corresponding to the digestion products of pUC19-Ku-LigD and cyclized P28 (Fig. S6), respectively. These results supported that Dt-Ku and Dt-LigD could function in *E. coli* to cyclize the linear DNA and both activities were needed. The electro-transformation of *E.coli* was 2.4 X 10^4^ CFU/μg DNA. The cyclization efficiency was calculated as 228 CFU/μg DNA in *E. coli* DH5α expressing Dt-Ku and Dt-LigD. The ratio of Amp/Kan resistant colonies was about 100:1. All the results were performed in triplicates.

### Recombination of linear DNA in *Dietzia* sp. DQ12-45-1b via NHEJ pathway

Since linear DNA could be cyclized in strain DQ12-45-1b by NHEJ activity *in vivo*, we assumed that it could act as a non-replicable plasmid that could further be used to inactivate a target gene by homologous recombination. To verify this hypothesis, we constructed a linear alk-Km DNA fragment containing a mono homologous arm DNA fragment of the *alkW1* gene and the kanamycin resistance cassette. After transforming 200 ng of this linear alk-Km DNA fragment into *Dietzia* sp. DQ12-45-1b, colonies were recovered in the presence of kanamycin. Among them, eight colonies were randomly selected and subjected to PCR amplification with primers alkW1-F and alkW1-R to amplify the *alkW1* gene. DNA fragments with the expected size (about 3.6 kb) were amplified from four out of the eight colonies, indicating that half of the colonies presented the correct insertion into the *alkW1* gene (Fig. 2A). Sequencing of these DNA fragments further confirmed the successful insertion of linear DNA into the *alkW1* gene in the genome (Fig. 2B). The recombinant efficiency was approximately 3×10^-6^.

**FIG 2.**
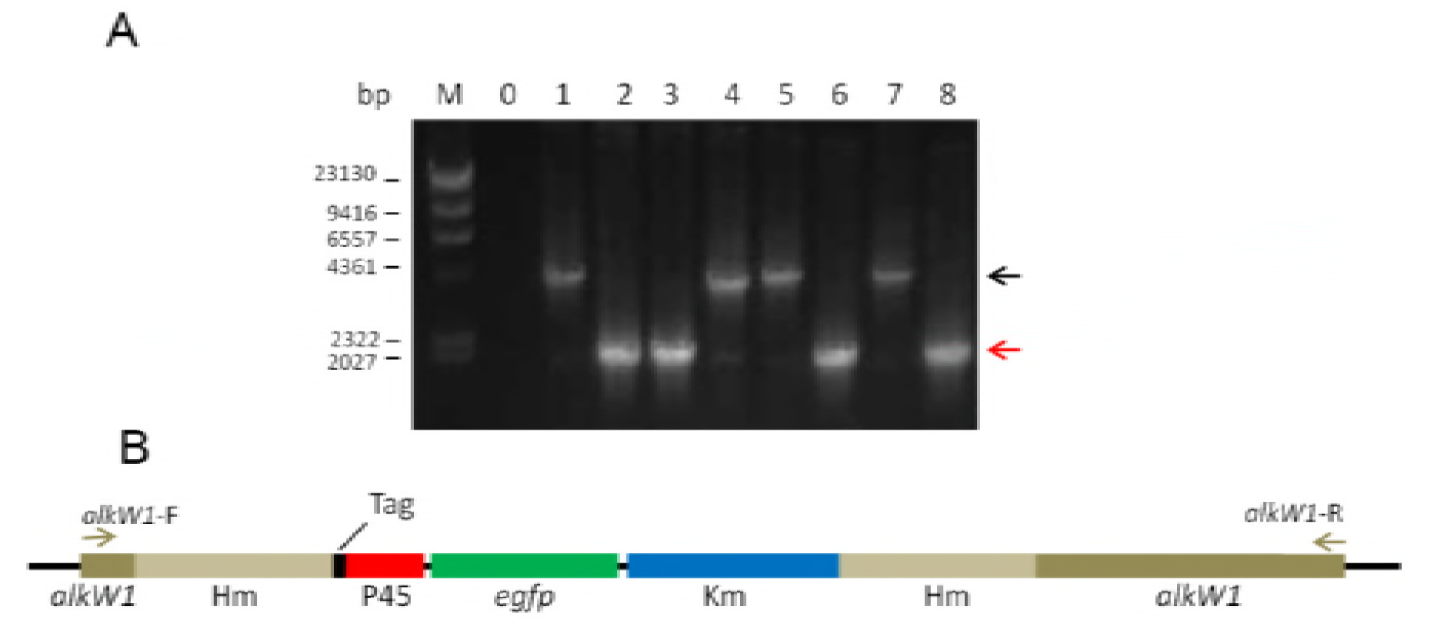
Identification of *alkW1* mutant in *Dietzia* sp. DQ12-45-1b by mono-homologous linear DNA. A, Ethidium-bromide-stained agarose gel showing typical results of PCR identification of eight colonies. M, DNA marker; 0, blank control. 1–8, different colonies. The bands designated with the black arrow are the *alkW1* mutant colonies with correct size. The bands designated with the red arrow are the unchanged *alkW1* colonies. B, Schematic diagram showing the linear DNA recombination pattern. P45, promoter p45 (119 bp); Hm, mono-homologous arm (410 bp); Km, kanamycine (795 bp).

To further verify the gene insertion, we carried out western blotting with the proteomes of *Dietzia* sp. DQ12-45-1b and *alkW1*^-^, which were grown in GPY medium or mineral medium with hexadecane as the sole carbon source. AlkW1 was induced in *Dietzia* sp. DQ12-45-1b by hexadecane, but not detected in the *alkW1*^-^ mutant strain either cultured in GPY or with hexadecane (Fig. 3A). As expected, the growth of the *alkW1*^-^ mutant strain in hexadecane as the sole carbon source was inhibited within eight days of incubation (Fig. 3B). These results proved that *alkW1* gene was successfully inactivated by the mono-homologous arm DNA fragment integration. Using the mono-homologous linear DNA recombination, we also successfully disrupt nine genes of *Dietzia* sp. DQ12-45-1b (Table 2), as confirmed by the PCR analysis (Fig. S7), the presence of green fluorescence (Fig. S8), and sequencing (Supplementary file 1). The recombinant efficiencies were approximately about 1.2 × 10^-6^-6.5 × 10^-5^ (Table 2). Besides, the successful inactivation of another gene *alkX*, the regulator of *alkW1* using this method (38), suggested the reliability of this method. In addition, the mutant was continuously cultured for 30 re-inoculations in GPY medium without antibiotics with no change in its genotype (data not shown), which demonstrated that the insertion was stable and heritable.

**FIG 3.**
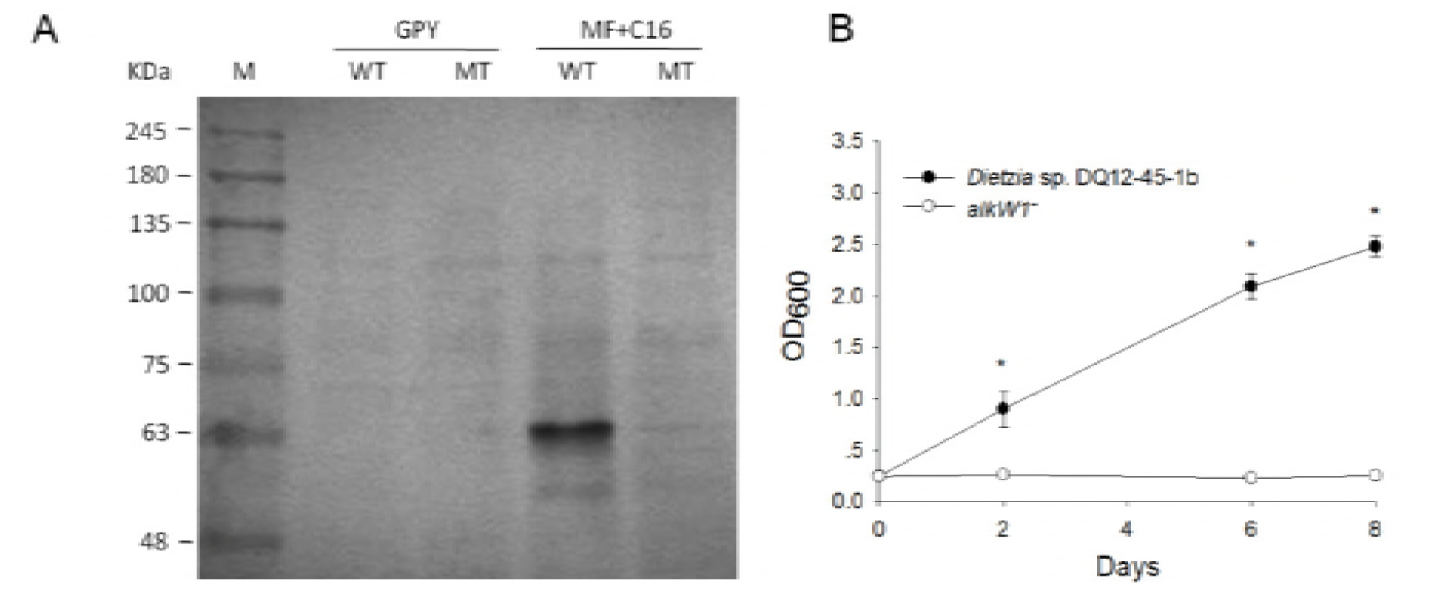
Functional characterization of the strain *alkW1*^-^. A, Western blot analysis of the AlkW1 protein expression in *Dietzia* sp. DQ12-45-1b and the strain *alkW1*^-^. Cells grew in GPY as negative control; WT cells grew in MF+ C16 as positive control; WT, wild type *Dietzia* sp. DQ12-45-1b; MT, the strain*alkW1*^-^; GPY, cells grown in GPY medium; MF+C16, cells grown in minimal medium containing hexadecane. B, Growth curve of strain DQ12-45-1b and strain *alkW1*^-^ in minimal medium with hexadecane as the sole carbon source.

### Distribution of Ku and LigD in prokaryotic organisms

By searching the *ku* and *ligD* homologous genes in the available 25,270 bacterial and 528 archaeal genomes using KO annotation, we identified 6,118 *ku* homologous genes in 5,098 bacterial and 13 archaeal genomes belonging to 14 bacterial phyla and Euryarchaeota. We also identified 18,952 *ligD* homologous genes in 7,631 bacterial and 64 archaeal genomes belonging to 26 bacterial phyla and 6 archaeal phyla (Fig. 4). Among them, 4,783 bacterial and 13 archaeal genomes contained both genes. Strikingly, about 76% and 77% of the total Actinobacteria genomes analyzed contained *ku* and *ligD* homologous genes, respectively, indicating that NHEJ might be common in Actinobacteria. Phylogenetic analysis showed that *ku* genes could be clustered into six clusters (Fig. 5), largely matching their taxonomic classification, suggesting the conservative property of the *ku* genes. In contrast, the *ligD* genes could be clustered into five clusters and their phylogenetic tree topology was much different from their taxonomic classification, suggesting possible vigorous gene transfer of *ligD* genes among microorganisms (Fig. 5).

**FIG 4.**
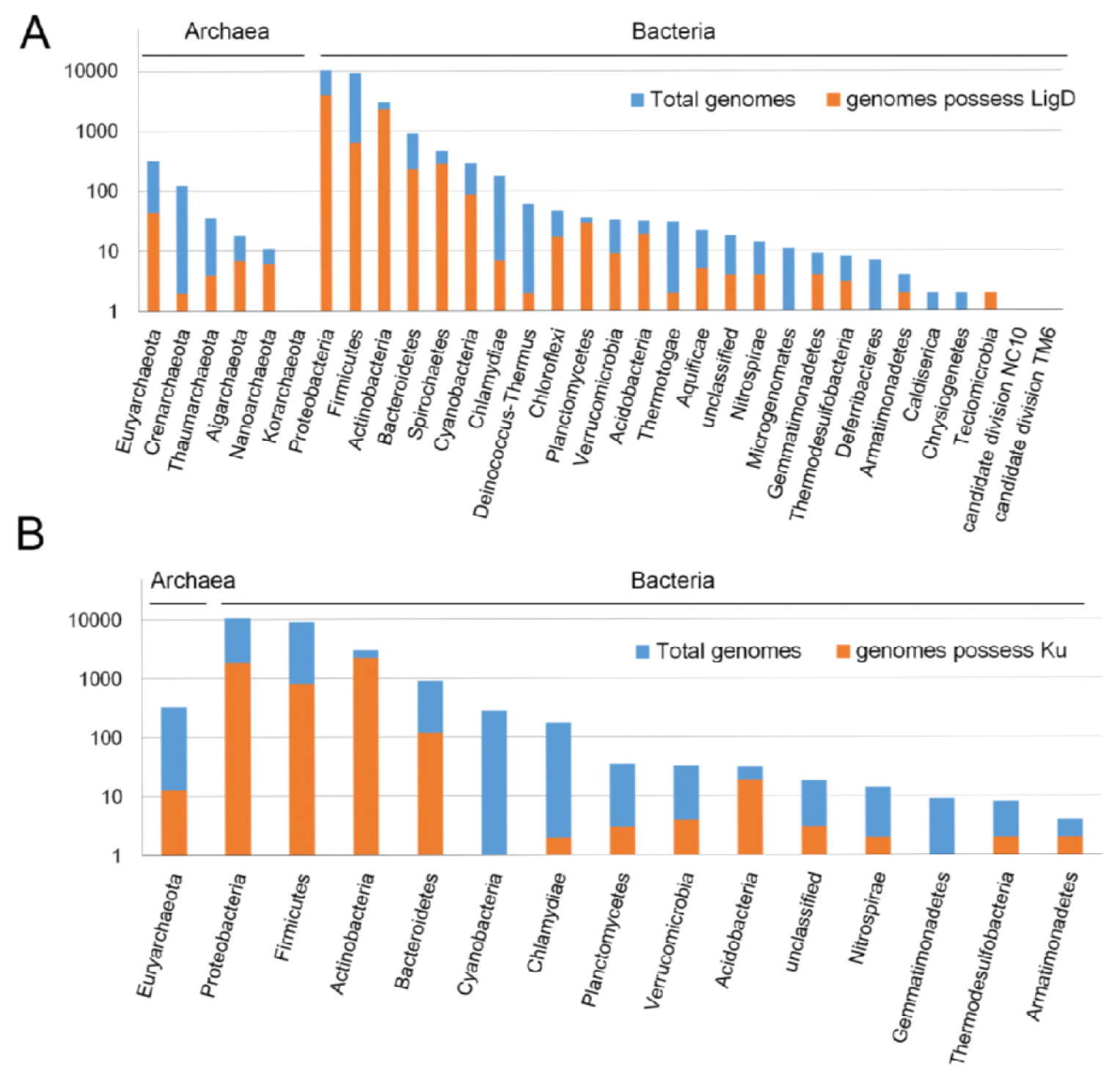
Distribution of *ku* and *ligD* homologous genes in microbial genomes at the phylum level. A, Distribution of *ligD* in microbial genomes at the phylum level; B, Distribution of *ku* in microbial genomes at the phylum level.

**FIG 5.**
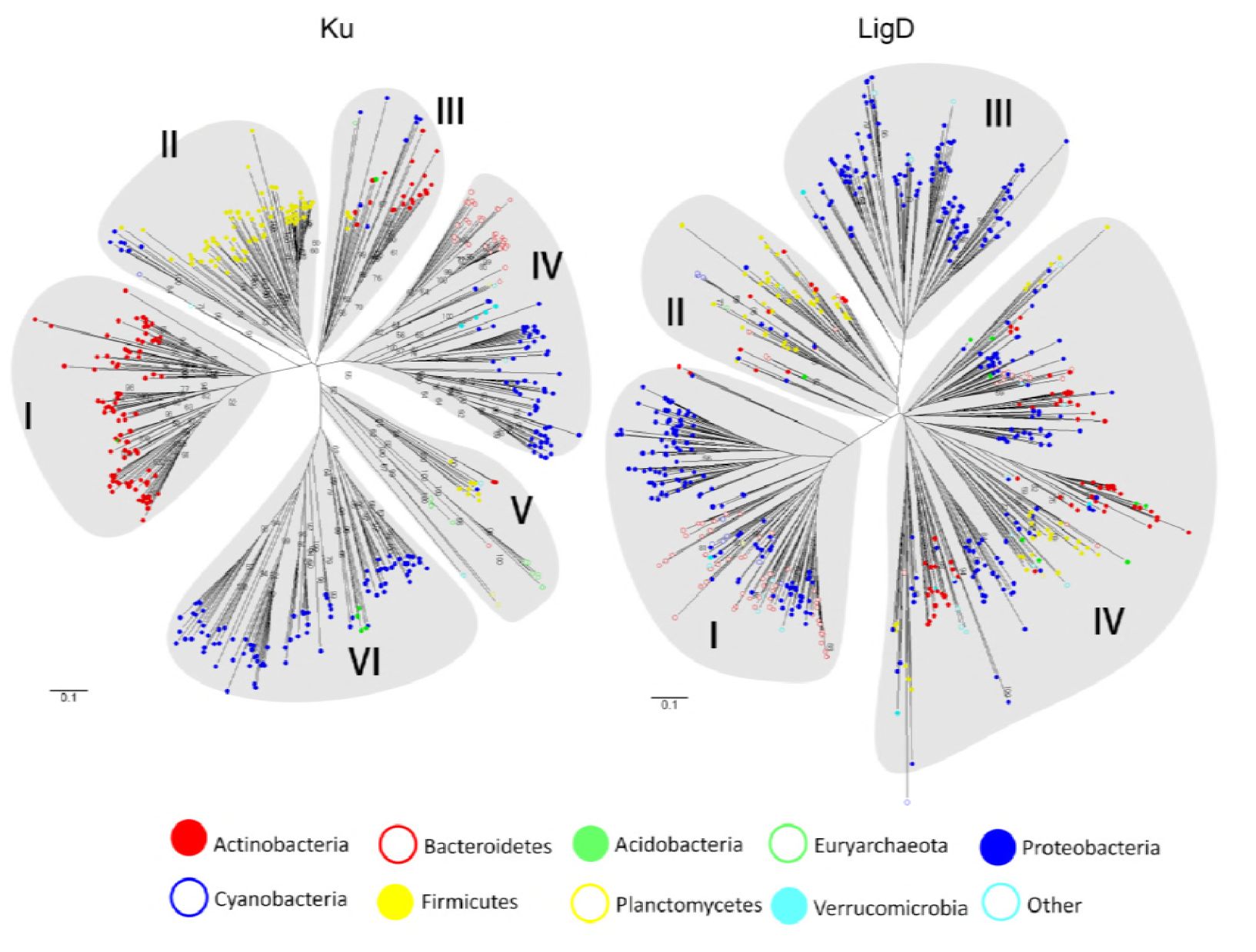
Phylogenetic relationships of Ku and LigD based on amino acid sequences in prokaryotic organisms. The phylogenetic trees were constructed using the neighbor-joining method in MEGA6 (45). The trees were bootstrapped with 500 replicates, indicated at the respective nodes.

The *ku* genes were generally juxtaposed with *ligD* genes (Fig. 6A), for which a functional association between Ku and LigD was suggested (21). In contrast, the Dt-Ku and Dt-LigD coding genes were not adjacent in *Dietzia* sp. DQ12-45-1b, but within 9.8 kb to each other (Fig. 6A). Similar gene arrangements were detected in other *Dietzia* species such as *Dietzia alimentaria*72, *Dietzia cinnamea* P4, and *Dietzia* sp. UCD-THP (Fig. S9A), whose Ku shared 72–81% amino acid identity with Dt-Ku. All the Ku protein sequences retrieved had a Ku core functional domain, which could bind DNA ends and transiently bring them together (Fig. 6B), suggesting the Ku functions in these strains (21). All the LigD sequences presented an ATP-dependent DNA ligase domain (Fig. 6C), which was essential in NHEJ (13, 14). The Ku core and ATP-dependent DNA ligase domain found in all Ku and LigD proteins suggested the intact NHEJ functions in all these strains containing both *ku* and *ligD.* Besides, Dt-LigD and its homologs from other *Dietzia* species contained a polymerase domain and a phosphoesterase domain from the N-terminal to the C-terminal, which was similar to that of the homolog Mt-LigD in *Mycobacterium* (Fig. 6C and Fig. S9B).

**FIG 6.**
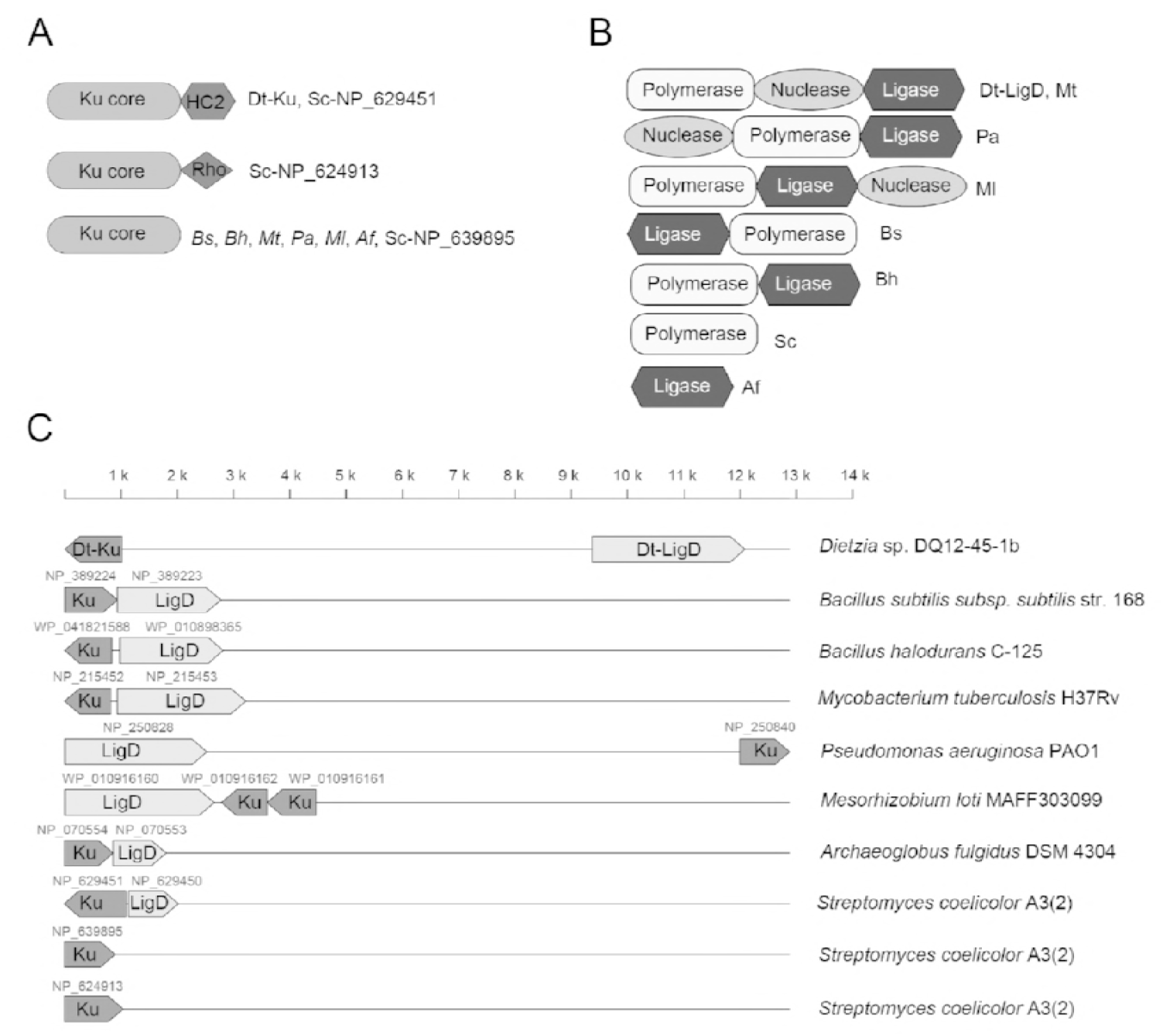
Organization of genes encoding Dt-Ku and Dt-LigD and their protein domain architecture. A, Protein domain architecture of Ku. Ku core, the Ku core domain; HC2, histone H1-like nucleoprotein domain; Rho, helix–extension–helix domain from the bacterial transcription termination factor Rho. B, Protein domain architecture of LigD. C, Gene organization of *ku* and *ligD* operons. The direction of the arrow indicates the direction of the transcription. The numbers above the arrows are the GenBank accession IDs of their corresponding genes. Sc, *Streptomyces coelicolor*; Bs, *Bacillus subtilis*; Bh, *Bacillus halodurans*; Mt, *Mycobacterium tuberculosis*; Pa, *Pseudomonas aeruginosa*; Ml, *Mesorhizobium loti*; Af, *Archaeoglobusfulgidus*.

## DISCUSSION

As the major DSB repair pathway, HR has been extensively studied in *E. coli* and widely used for gene manipulation. This classical genetic engineering method uses incompatible or suicide plasmids and requires a time-consuming process. A phage-mediated HR, usually referred to as “recombineering”, constructs linear DNA with short homologies (52) and was conveniently applied in genomic manipulation of *E. coli*, *Mycobacterium*, *Salmonella*, and *Shigella* (47, 53–55). Unfortunately, the known phage systems are not applicable for recombineering many other bacteria of interest. For example, 666 novel species were published in 2013 (LPSN-list of prokaryotic names with standing in nomenclature, http://www.bacterio.net/), a lot of which are of great importance in bioremediation, chemical production, and human health. However, it is barely possible to find compatible phage systems as well as shuttle vectors for these strains. Although ZFNs, TALENs, and CRISPR have been developed for precise genome editing in eukaryotes (8–10), they also need shuttle vectors to express a nuclease in the host strain. The lack of shuttle vectors expressing these systems is additionally hindering their use for diverse prokaryotes. In this study, we proposed a novel and much simpler gene inactivation method by using the NHEJ pathway mediated by Ku and LigD proteins in *Dietzia* sp. DQ12-45-1b without additional vectors or genome editing systems.

After the linear DNA with mono-homology was transformed into *Dietzia* sp. DQ12-45-1b cells, it performed self-cyclization *in vivo*, recombined with the target homologous sequence in the genome by HR and produced the mutant strain (Fig. 7) (38). Although the efficiency was not very high, around 10^-5^ - 10^-6^, when targeting the genes in *Dietzia* sp. DQ12-45-1b, it may be acceptable for those newly discovered and important bacteria without compatible genetic manipulation system. Although NHEJ was firstly found in mammalian cells and thought to be specific to eukaryotic organisms, a number of studies showed that NHEJ also exists in prokaryotic organisms (15, 16, 18–23, 56). Ku were proved to be essential in NHEJ in the strain DQ12-45-1b. It was interesting that we tried to disrupt the *ku* and *ligD* genes using two methods, the mono-homologous recombination method as described in this paper and the double-homologous recombination method described previously (47), but neither *ku* nor *ligD* mutants was obtained using the mono-homologous linear DNA recombination method. In contrast, *ku* gene could be replaced using double-homologous recombination method which expressed additional recombination enzymes. The result suggested unknown roles of Ku in the recombination process using mono-homologous linear DNA. It was notable that no *ligD* mutant was obtained using both two methods. LigD, as the main ligase of NHEJ, could repair DSBs that had a lethal effect on cell mitosis unless repaired in time (57–60). According to previous reports, *ligD* could be knocked out from *Mycobacterium smegmatis*, because there was another ATP-dependent DNA ligase (LigC) that could provide a backup function of LigD-independent error-prone repair of blunt-end DSBs in the strain (61). However, in *Dietzia* sp. DQ12-45-1b, only one *ligD* was identified, and its disruption might be lethal. Furthermore, there was no cyclization of linear plasmid DNA in *ku* mutant of strain DQ12-45-1b, implying that *ku* might be critical in the process of linear DNA cyclization.

**FIG 7.**
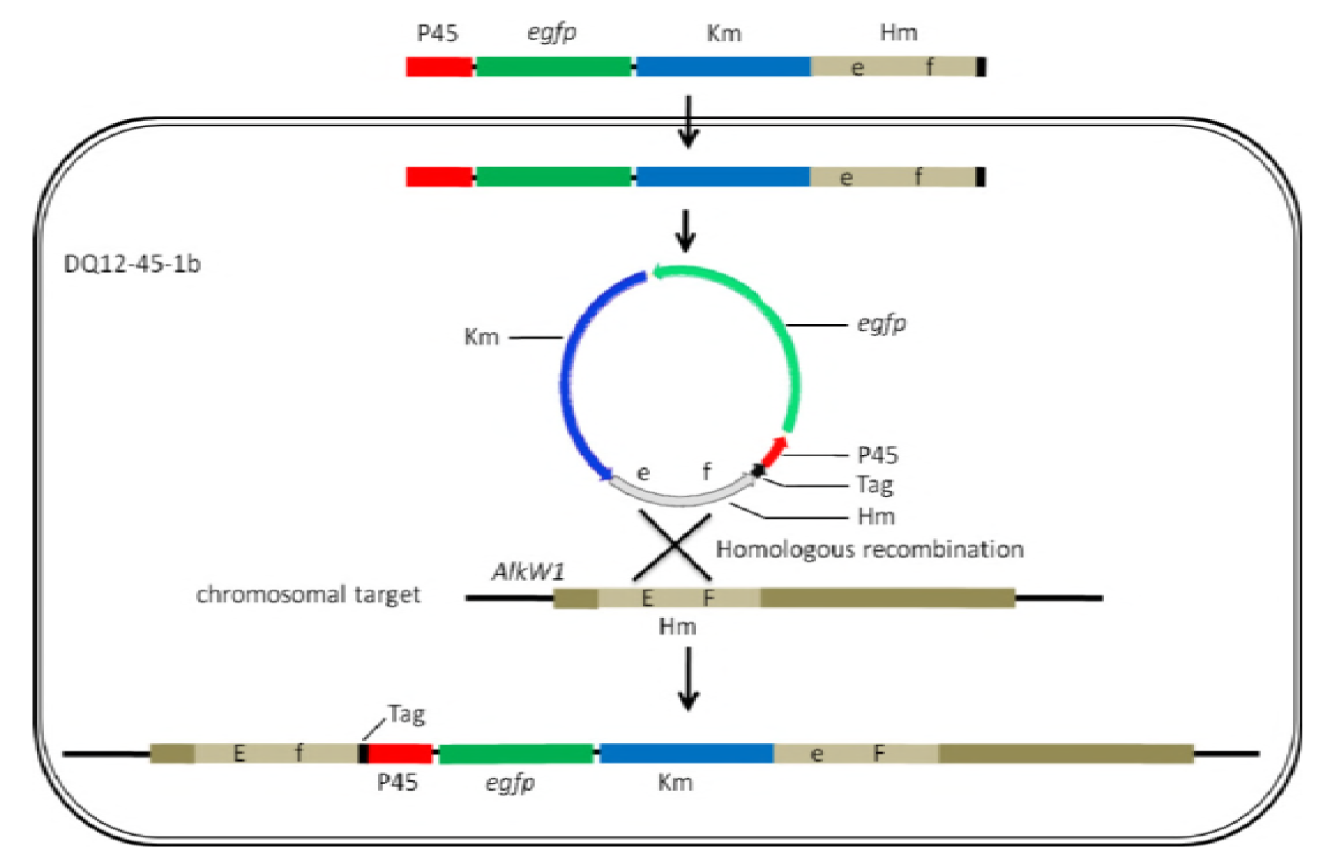
Mechanism of mono-homologous linear DNA for target gene mutant. The linear DNA transformed into bacterial cells is self-cyclized and then recombined with the chromosomal gene.

In addition, 5,111 and 7,695 out of the 25,798 bacterial and archaeal genomes available in the IMG database contain *ku* and *ligD* homologous genes, respectively. The distribution of *ku* and *ligD* genes may suggest the wide distribution of the NHEJ pathway in microorganisms.

Consequently, the introduction of mutagenesis during the cyclization might have played an important role in the genome diversification, which could increase the natural microbial diversity and adaptability. In addition, the wide distribution of *ku* and *ligD* genes may also suggest that the mono-homologous linear DNA recombination by using the NHEJ pathway could be commonly used as a gene inactivation method for diverse prokaryotic organisms, especially for newly discovered, but important microorganisms. Especially, more than 3/4 of the genomes of Actinobacteria, with high GC content and not easy for genetic manipulation, express the *ku* and *ligD* genes, suggesting that the mono-homologous linear DNA recombination is a potentially good method, at least for Actinobacteria. However, more studies are needed to test the versatility of this method.

For organisms lacking Ku and LigD, the mono-homologous linear DNA recombination method requires the construction of a compatible and inducible Ku and LigD system, i.e., one plasmid needs to be constructed to express Ku and LigD. In this case, the expression of Ku and LigD could be controlled by appropriate inducible promoters as described for the expression of Mt-Ku and Mt-LigD in *E. coli* (17). Once the plasmid expressing Ku and LigD is constructed, it can be used for disrupting any gene without constructing new plasmids. Thus, this method is still simpler than the classic approaches requiring incompatible or suicide plasmids.

This method presents two major concerns. One is how to improve the cyclization efficiency and the other is how to reduce the risk of linear DNA degradation by nucleases *in vivo.* In prokaryotic organisms, NHEJ activity was particularly efficient during the stationary phase to counteract DSBs induced by heat, desiccation, and other factors (62). However, the electroporation efficiency reached its highest value when the recipient cells were in the early exponential phase (33). A balance is therefore necessary to increase the cyclization and recombination activities and to increase the transforming efficiency. Several methods have been developed, including heating the cells, to increase both the transforming efficiencies and the NHEJ process (33), which may promote the efficiency of this method.

Unlike in yeast, linear dsDNA in bacteria such as *E. coli* can be rapidly degraded by nucleases (26). This phenomenon was also observed in strain DQ12-45-1b during sequence deletions when cyclizing the linear DNA LA18, HD18, and HB18 *in vivo* (Fig. 1). In *E. coli*, *recB*, and *recC* are the major nucleases degrading the dsDNA and are essential for the normal activities of the bacterium. Deletion of *recB* and *recC* resulted in poor growth of *E. coli*, producing up to 80% of nonviable cells (12). To inhibit the nuclease activities and promote the recombination efficiency, several phage systems such as RecET system and λ Red system were developed, which again require plasmids (53, 63). For example, in the λ Red system, the *exo*, *bet*, and *gam* genes are under the *lac* promoter control on a multicopy plasmid. Gam function could inhibit RecBCD nuclease to protect the transformed linear DNA from being degraded. Then, with the help of Exo and Bet, gene recombination was conducted (12). Another trial was to recombine linear DNA in wild-type *E.coli* containing the RecBCD nuclease (64). In this case, special sites were engineered in the linear DNA that decreased the RecBCD activities (64). Similarly, when we added a triplet cytidine (CCC) tag at one terminus of the linear DNA (LA18) by PCR, the long deletions were unidirectional and occurred at the opposite terminus with lower “G+C” contents (Fig. 1), suggesting that addition of the CCC tag could protect the target sequence from being digested by nucleases and increased the recombination efficiency. Additionally, electroporation itself has been suggested to reduce DNA degradation by RecBCD nuclease and to allow recombination with linear dsDNA (65).

If this method is reliable, it could be used not only to inactivate one gene by inserting the selective cassette into the chromosome (Fig. S10A), but also for gene over-expression and exogenous gene insertion. For example, for over-expression of a gene, the cassette consists of a selectable integrated gene that would be used as the mono-homologous DNA (Fig. S10B). For exogenous DNA insertion, the cassette could be linked to an exogenous DNA before the homologous arm (Fig. S10C).

In conclusion, we proposed an easy and fast method for gene inactivation in *Dietzia* sp. DQ12-45-1b, using a linear DNA with mono-homology. This method uses the NHEJ pathway and it might be applicable to prokaryotic organisms that harbor genes encoding Ku and LigD, especially for the newly identified and non-model species. Because the linear DNA with cassettes and homologous DNA could be amplified by fusion PCR conventionally, these procedures should be rapid, convenient, and reliable.

## ACKNOWLEDGMENTS

We thank Dr. Guang Hu for generously providing the antibody of polyclonal mouse anti-AlkW1, Ms. Hui Fang for her kindly help in bioinformatic analysis, Ms. Xiao-Yu Qin for her help in Ku knockout, and we thank Dr. Yue-Qin Tang for reading the manuscript and discussions.

## Footnotes

1 The authors wish it to be known that, in their opinion, the first two authors should be regarded as joint First Authors

